# Unravelling cancer subtype-specific driver genes in single-cell transcriptomics data with CSDGI

**DOI:** 10.1101/2023.08.23.554393

**Authors:** Meng Huang, Jiangtao Ma, Guangqi An, Xiucai Ye

## Abstract

Cancer is known as a heterogeneous disease. <u>C</u>ancer <u>d</u>river <u>g</u>enes (CDGs) need to be inferred for understanding tumor heterogeneity in cancer. However, the existing computational methods have identified many common CDGs. A key challenge exploring cancer progression is to infer cancer subtype-specific driver genes (CSDGs), which provides guidane for the diagnosis, treatment and prognosis of cancer. The significant advancements in single-cell RNA-sequencing (scRNA-seq) technologies have opened up new possibilities for studying human cancers at the individual cell level. In this study, we develop a novel unsupervised method, *CSDGI* (<u>C</u>ancer <u>S</u>ubtype-specific <u>D</u>river <u>G</u>ene <u>I</u>nference), which applies Encoder-Decoder-Framework consisting of low-rank residual neural networks to inferring driver genes corresponding to potential cancer subtypes at single-cell level. To infer CSDGs, we apply *CSDGI* to the tumor single-cell transcriptomics data. To filter the redundant genes before driver gene inference, we perform the differential expression genes (DEGs). The experimental results demonstrate *CSDGI* is effective to infer driver genes that are cancer subtype-specific. Functional and disease enrichment analysis shows these inferred CSDGs indicate the key biological processes and disease pathways. *CSDGI* is the first method to explore cancer driver genes at the cancer subtype level. We believe that it can be a useful method to understand the mechanisms of cell transformation driving tumours.

**Author summary:** Cancer is recognized as a complex disease with diverse characteristics. In order to comprehend the diversity within tumors, it is essential to infer cancer subtype-specific driver genes (CSDGs), which offer valuable insights for investigating cancer progression and treatment. The remarkable progress made in single-cell RNA-sequencing (scRNA-seq) technologies has ushered in new prospects for studying human cancers at the cellular level. Cancer Subtype-specific Driver Gene Inference (CSDGI) is a novel unsupervised method proposed. In our study, we use Encoder-Decoder-Framework to infer driver genes specific to cancer subtypes in the CSDGI. We apply CSDGI to three tumor single-cell transcriptomics data. The experimental results have shown the effectiveness of CSDGI. Furthermore, functional and disease enrichment analyses illustrate that these inferred CSDGs shed light on crucial biological processes and disease pathways. Our collection of driver genes will serve as a valuable resource in unraveling the mechanisms driving cell transformation in tumors.

## Introduction

Cancer is a heterogeneous disease characterized by the unstable cellular growth [1, 2], which is caused by a set of genes. These genes can drive tumorigenesis as drivers, and thereby are known as cancer driver genes (CDGs) [1]. They may affect the homeostatic development for a collection of critical cellular function containing cell proliferation, cell differentiation, cell death, etc. To advance the research of tumor emergence and evolution, it is crucial to infer CDGs [3, 4]. Further, cancer molecular subtypes play a crucial role in deepening our insights of cancer within tumor cell subsets as representing a heterogeneous disease instead of a single disease [5], which may suggest therapeutic opportunities. Therefore, the discovery of these CDGs across cancer subtypes helps explore cancer progression. To guide the diagnosis, treatment and prognosis of cancer, it is also necessary to infer cancer subtype-specific driver genes (CSDGs).

Previous studies have shown that CDGs may be tissue-specific [6] and condition-specific [7, 8]. For example, CDGs can be identified in a large number of tumor samples across cancer subtypes using bulk transcriptome data by RNA sequencing (RNA-seq) from The Cancer Genome Atlas (TCGA) consortium [6, 9]. To infer the driver gene in large cohorts, many projects have been performed in the whole genome of thousands of tumour samples under the International Cancer Genome Consortium (ICGC) [10, 11]. Besides, transcriptomic studies have supported most cancer subtype discoveries rather than other omics data [12]. Nevertheless, previous computational methods [13–16] using bulking transcriptome data mostly identify CDGs involved in all of cells, rather than CDGs specific to cell subpopulations (cancer subtypes). This may conceal the heterogeneity of CDGs across different cancer subtypes. Fortunately, more transcriptome studies have become possible due to the advancements in single-cell RNA-sequencing (scRNA-seq) technologies [17–20]. In contrast to traditional bulk RNA-seq providing average gene expression of cell groups [21–22], scRNA-seq allows for the quantification of gene expression in individual cells, which facilitates the analysis of cellular differences [24–26]. This is particularly valuable for cancer-related research, as scRNA-seq allows for the identification and analysis of intratumoral heterogeneity [27–29]. As a result, researchers can obtain an in-depth understanding of cancer subtypes, cellular populations, and the functions of individual cells within tumor samples at single cell level [25, 30–31]. Accordingly, the scRNA-seq can provide a way to infer CDGs at the single-cell or cancer-subtype level.

By using bulk transcriptome and other omic data, several methods have been developed to identify CDGs. For example, Akavia et al. developed CONEXIC to unveil potential CDGs in the deleted tumor region by combining gene expression data with copy number change [13]. Ng et al. proposed PARADIGM-SHIFT to infer gene activity utilizing pathway-level information, gene expression and copy number, which helps predict mutation drivers in cancer processes [14]. Paull et al. developed TieDIE to find small cancer driver pathway by using genomic and transcriptomic perturbations, which may predict transcription factor in cancer [15]. Chen et al. presented MAXDRIVER to infer CDGs by integrating genomic data and heterogeneous networks [16]. However, the above computational methods are limited to unveil CDGs specific to tissues or samples since they do not take into account the relationships between genes and cancer subtypes. To understand the heterogeneity of cancer across different cancer subtypes, it is necessary to infer cancer subtype-specific driver genes.

In ths work, we present a novel unsupervised computational method, *CSDGI* (<u>C</u>ancer <u>S</u>ubtype-specific <u>D</u>river <u>G</u>ene <u>I</u>nference) to infer driver genes specific to potential cancer subtypes at single-cell level. By only using single-cell transcriptomics data rather than a large cohort, the proposed *CSDGI* method applies the Encoder-Decoder-Framework that consists of low-rank residual neural network to selecting genes that are most associated with the identified cancer subtypes. *CSDGI* ranks potential driver genes based on the association between genes and cancer subtypes. In each cancer subtype identification module, genes with higher rankings are more likely to be drivers specific to the current cancer subtype. To filter the redundant genes, the differential expression genes (DEGs) are performed before inferring driver gene. We apply *CSDGI* to the real tumor single-cell transcriptomics datasets, which infers CSDGs relating to cancer subtypes. The experimental results demonstrate that *CSDGI* can be an effective method to infer driver genes for identified cancer subtypes. Especially, the associated driver genes can be applied to explore the interpretable biological meaning of each cancer subtype. Systematic gene ontology and disease enrichment analysis demonstrates the potential functions and key biological processes of identified CSDGs. *CSDGI* is the first method to explore CSDGs at the cancer subtype level. These CSDGs and the proposed methodology provide a new way to better understand the mechanisms of tumorigenesis, help classify cancer subtypes and explore the genotypic status of tumors. We believe that *CSDGI* can be a useful method to understand the mechanisms of cell transformation driving tumours, and cancer progression.

## Materials

### Single-cell transcriptomics data in the human cancer

In this study, we use three single-cell transcriptomics datasets as given by the data repository NCBI Gene Expression Omnibus. We download melanoma, breast cancer, and chronic myeloid leukemia gene expression data from GSE72056 [29], GSE75688 [30] and GSE76312 [31]. The summary of these datasets is provided in Table 1. In the melanoma dataset (GSE72056), these cells comprise 1257 malignant melanoma tumor cells and 3388 benign tumor cells. Besides, the breast cancer dataset (GSE75688) consists of 317 tumor cells and 198 nontumor cells. Both GSE72056 and GSE75688 datasets were transformed using log𝑇𝑃𝑀 as inputs for the model. Additionally, the chronic myeloid leukemia dataset (GSE76312) includes cells from the human chronic myeloid leukemia, comprising 902 tumor cells and 232 normal cells. The dataset from GSE76312 was transformed by log𝑅𝑃𝐾𝑀 as inputs of model.

**Table 1.**
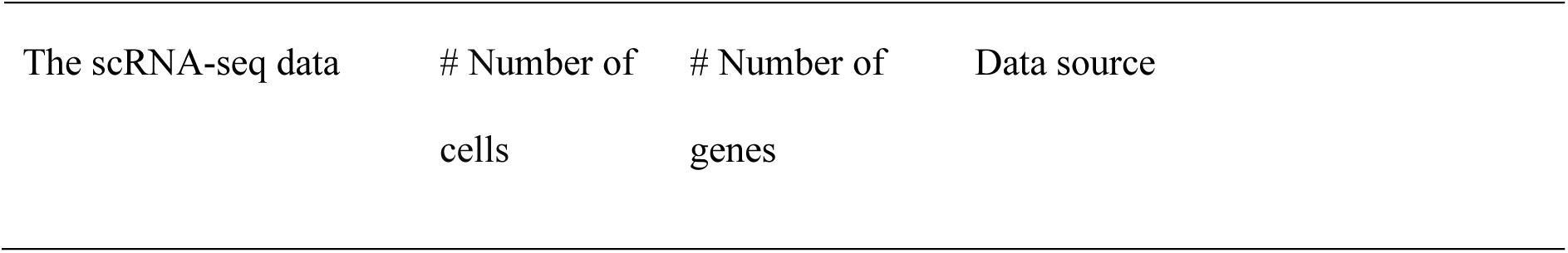
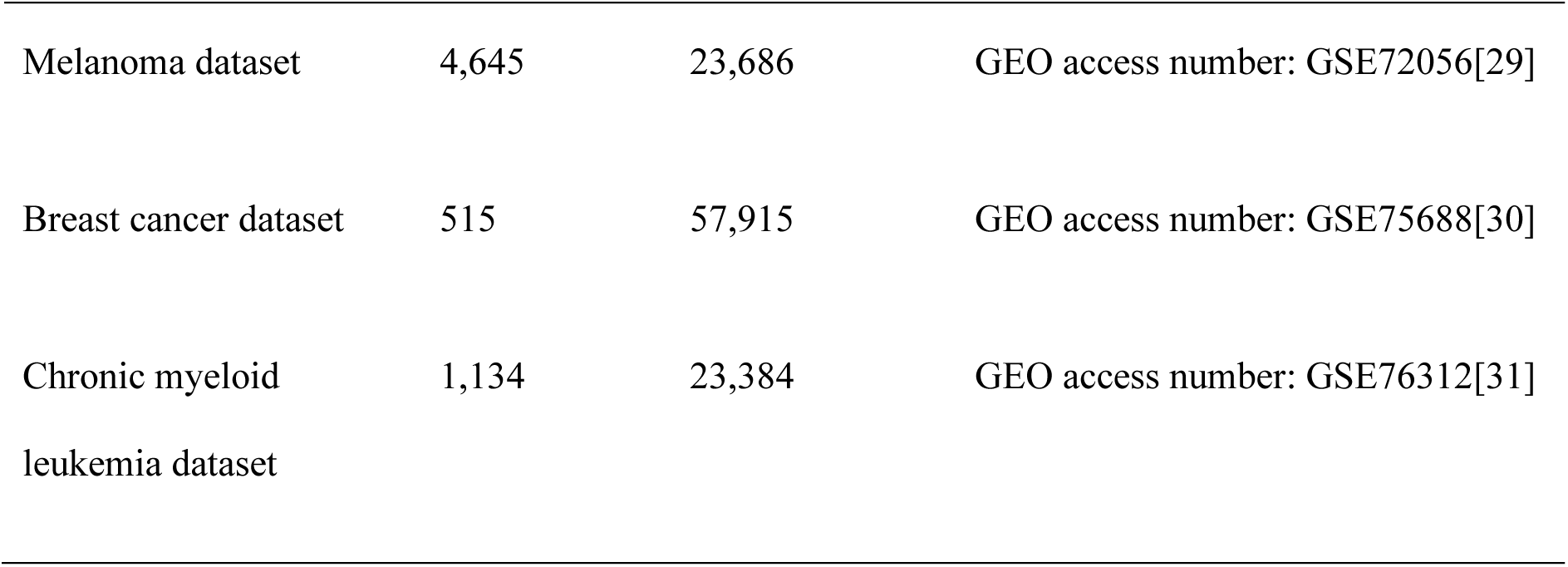
The summary of single-cell transcriptomics data of human cancer used in this study.

| The scRNA-seq data | # Number of<br>cells | # Number of<br>genes | Data source |
| --- | --- | --- | --- |
| Melanoma dataset | 4,645 | 23,686 | GEO access number: GSE72056[29] |
| Breast cancer dataset | 515 | 57,915 | GEO access number: GSE75688[30] |
| Chronic myeloid<br>leukemia dataset | 1,134 | 23,384 | GEO access number: GSE76312[31] |

### Data pre-processing

To investigate the intrinsic transcriptomic signatures of tumor cells, we apply a filtering process to the gene expression data. We filter out genes that are expressed in less than 𝑡% of cells, where 𝑡% is set to 6% based on the previous study [32]. Additionally, we also filter out genes that are expressed in more than (100 − 𝑡)% of cells, as these ubiquitous genes are not helpful in inferring cancer subtypes. Following the gene filtering, we select the most 𝑟% variable genes by controlling the relationship between mean expression and variability. After gene filtering, there are 12693 genes in the melanoma dataset, 15451 genes in the breast cancer dataset, and 10862 genes in the chronic myeloid leukemia dataset. Subsequently, we utilize a R tool (EMDomics) [33] to obtain differential expression genes (DEGs) between tumor cells and benign (nontumor or normal) cells. As a result, we obtain 820 DEGs in the breast cancer dataset, 1048 DEGs in the chronic myeloid leukemia dataset, and 1170 DEGs in the melanoma dataset. These selected DEGs also help infer CSDGs in the breast tumor cells, the chronic myeloid leukemia tumor cells and the malignant melanoma tumor cells. More detailed information about DEGs can be found in Supplementary Material 1.

## Methods

### Proposed method overview

To identify cancer subtype-specific driver genes, we propose a novel unsupervised method, cancer subtype-specific driver gene inference (CSDGI) by using Encoder-Decoder-Framework. This workflow is shown in Fig 1. Firstly, we download real tumor scRNA-seq datasets including nontumor cells and tumor cells. These data are preprocessed to obtain the gene expression data 𝑋. Then, we apply Encoder-Decoder-Framework to infer driver genes for each cancer subtype. We rank genes according to their low-rank weights in the Encoder-Decoder-Framework, i.e., the importance of a gene in various cancer subtypes will result in its higher ranking. The output of the rank display the overall impact of each gene for every cancer subtypes. Finally, we perform downstream analysis with cancer subtype-specific driver genes.

**Fig 1.**
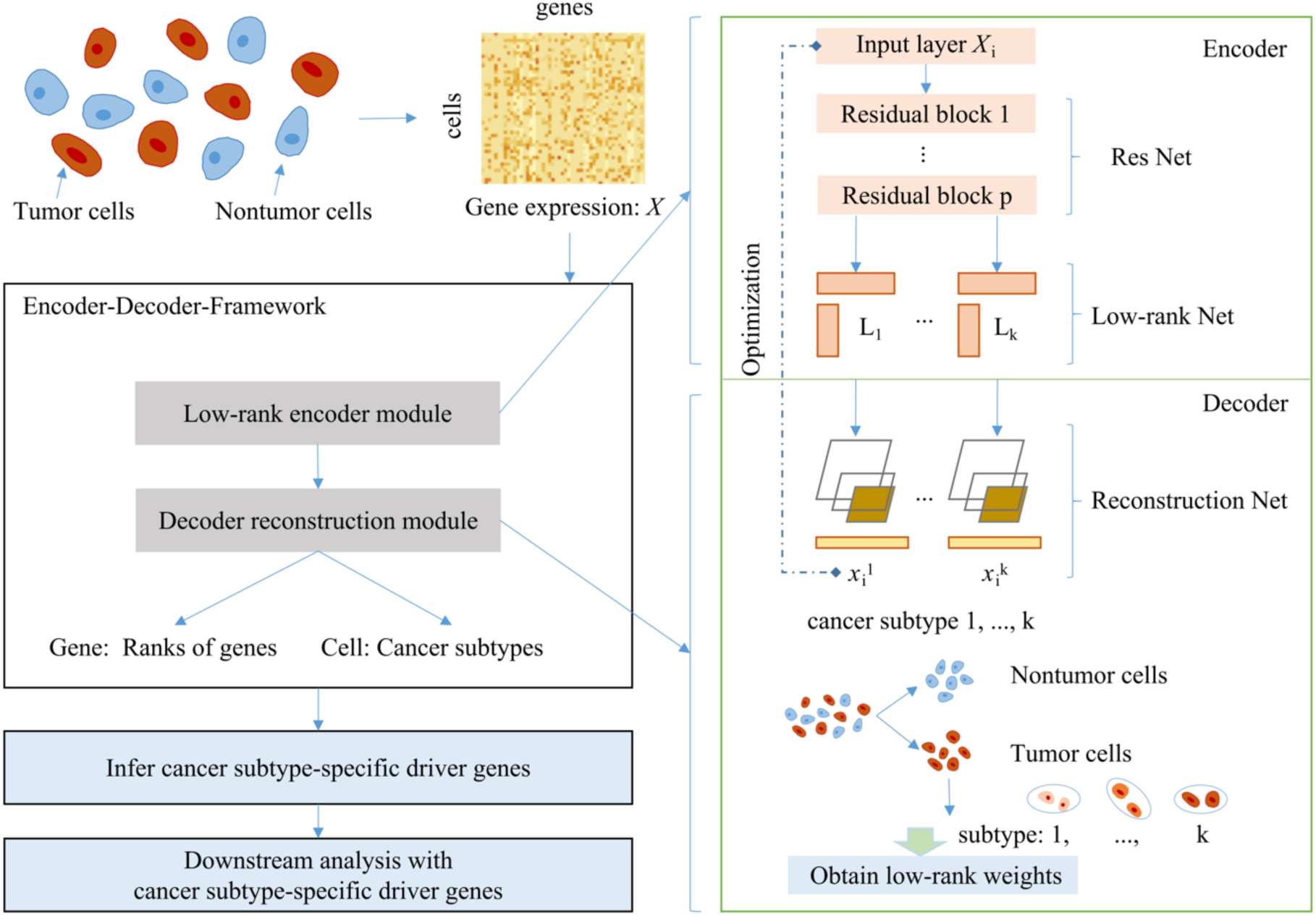
Workflow of CSDGI. After data pre-processing for real tumor scRNA-seq datasets, we infer different driver genes for different cancer subtypes using Encoder-Decoder-Framework and perform downstream analysis with cancer subtype-specific driver genes.

### Encoder-Decoder-Framework

Encoder-Decoder-Framework consists of low-rank encoder module and decoder reconstruction module. As shown in Fig 1, the single-cell gene expression matrix is represented as 𝑋 = [𝑋_1_, 𝑋_2_, 𝑋_3_,···, 𝑋_𝑚_]^𝑇^ ∈ ℝ^𝑚×𝑛^, where 𝑋_𝑖_ ∈ 𝑅^𝑛×1^ indicate the gene expression profile of the 𝑖-th cell. Here, 𝑚 and 𝑛 denote the number of cells and genes, respectively. 𝑘 is the number of estimated cancer subtypes In Encoder, motivated by the interpretable feature clustering method and the automatic association feature learning in scRNA data [34, 35], we apply the low-rank residual network to encoding the low-rank representation of each subtype. In detail, the gene expression of each cell 𝑋_𝑖_, 𝑖 ∈ {1, 2, . . ., 𝑚} is input to the input layer. The deeper residual neural network (ResNet) [36] aims to redefine the layers by acquiring an understanding of residual functions associated with the input layer. The learnable network parameters of ResNet represent the non-linearities between genes, which is shared across each subtype. The number of residual block is *p*. The objective function of ResNet is as follows:

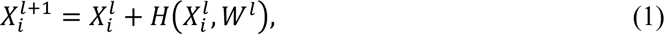

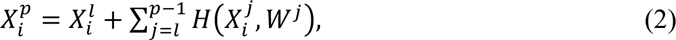

where H is the non-linear function representation, where the activate function is ReLU and the bias is 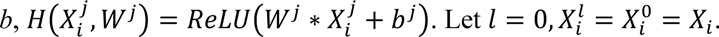 .

To represent the gene association across different genes in each cancer subtype, we utilize low-rank network that is performed using low-rank matrix [37, 38]. Here, we view *L*_k_ as the low-rank (r-rank) graph of cancer subtype *k*. The function of low-rank network is as follows:

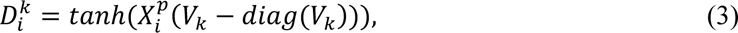

where 𝐿_𝑘_ = (𝐵_𝑘_)^-1^ 𝑉_𝑘_(𝑉_𝑘_)^𝑇^, 𝑉_𝑘_(𝑉_𝑘_)^𝑇^ is the *r*-rank matrix, 𝑉_𝑘_ ∈ ℝ^𝑚×𝑟^, the degree matrix of 𝑉_𝑘_(𝑉_𝑘_)^𝑇^ is 𝐵_𝑘_, the output is 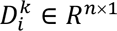. Empirically, we set *r* to 1.

In Decoder, we use the deep neural network to implement the Reconstruction Net. To obtain the reconstructed gene expressions 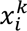, we perform the following calculations:

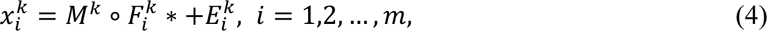

where 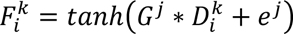, *G* and *e* are the weight and bias, ∘ is the dot product, *M* and *E* are the learnable weight matrix and bias for 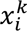 .

To minimize the loss between source gene value 𝑋_𝑖_ and reconstructed gene value 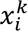, we apply the mean squared error (MSE) to them. The residual between 𝑋_𝑖_ and 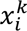 is calculated as follows:

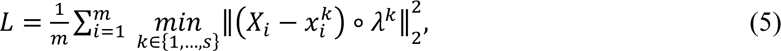

where *m* is the number of cells, *s* is the maximum number value for cancer subtype index *k*, the low-rank weight 𝜆*^k^* is the absolute value of 𝜇*_k_*, 𝜇*_k_* is the first nontrivial left eigenvector of (𝐼 − 𝐿*_k_*) of low-rank matrix in the Encoder. After obtaining optimal loss, the subtype index *k* with the minimum loss indicates a subtype label of each cell.

### Gene inference and downstream analysis

To infer CSDGs, we calculate the values of low-rank weight 𝜆*^k^* ∈ 𝑅^𝑛×1^ corresponding of each cancer subtype *k* after obtaining the optimal model of Encoder-Decoder-Framework. We also obtain the subtype label of each cell that is assigned according to the minimum loss of model. In each cancer subtype, each value of 𝜆*^k^* is sorted in descending order according to the weight values, where 𝜆*^k^* indicates the importances of genes in the cancer subtype *k*. We define the top 5% of genes as cancer subtype-specific driver genes in the sorted value of weight 𝜆*^k^*. In order to explore the biological meaning of CSDGs, we perform downstream analysis including subtype classification, functional and disease enrichment analysis.

### Measurement metrics

To quantitatively evaluate the performance of CSDGI, we adapt the accuracy metric for classification and adjusted rand index (ARI) for clustering. Theset metrics are defined as follows:

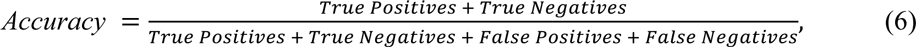

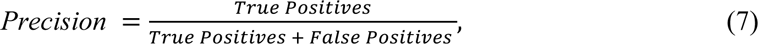

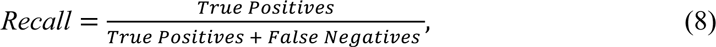

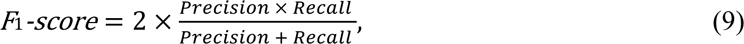

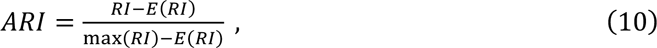

where True Positives represents the number of positive samples that are correctly classified as positive, True Negatives represents the number of negative samples that are correctly classified as negative, False Positives represents the number of negative samples that are incorrectly classified as positive, False Negatives represents the number of positive samples that are incorrectly classified as negative, RI is the rand index and ARI is the adjusted rand index to assess the cluster performance.

### Parameter settings

To identify unique driver genes distinguishing tumor cells from nontumor cells in three real tumor single-cell transcriptomics datasets, we set the value of 𝑘 to 2. To explore different driver genes associated with inferred cancer subtypes within tumor cells, the factoextra (R package tool) is used to determine the cluster number of tumor cells. Based on the results of the cluster number analysis, *k* is set to 2, 2, and 3 in the proposed CSDGI method for the breast tumor cells, the chronic myeloid leukemia tumor cells, and the malignant melanoma tumor cells, respectivcely. All model optimization experiments of CSDGI are iteratively conducted for 400 epochs using a single NVIDIA Tesla Ampere A100-PCIe-40GB GPU and the Ubuntu 18.04 system. Furthermore, all evaluation experiments are run independently 100 times and the average results are shown in the study.

## Results

### Differential expression genes analysis

To filter redundant genes, we use a R tool (EMDomics) to obtain DEGs between tumor cells and nontumor cells. Fig 2 presents different density plots of genes illustrating the distribution of various genes in three real tumor scRNA-seq datasets. Besides, we obtain all emd scores and *p*-value of preprocessing genes by using the EMDomics tool. We focus on genes having higher emd scores with *p*-value < 0.05. For example, the emd score results with melanoma dataset (GSE72056) are shown in Fig 2(A, B, C). Fig 2A demonstrates that the EMD score is 0.01 for the gene RP11-567G24.1, which represents the lowest value, while Fig 2C exhibits the highest EMD score of 23.9 for the gene TMSB4X in the breast cancer dataset. Fig 2B displays the density plot for the gene C18orf32, which shows 7.01 EMD scores. Notably, higher EMD scores indicate a greater distinction between tumor cells (group A) and nontumor cells (group B). Consequently, we select 820 genes with EMD scores ranging from 7 to 23.9 as DEGs in the breast cancer dataset. Similarly, Fig 2(D, E, F) show that we obtain 820 genes with emd scores varing from 2 to 13.57 as DEGs in the chronic myeloid leukemia dataset. As shown in Fig 2(G, H, I), 1170 genes with emd scores more than 7 are regarded as DEGs in the melanoma dataset. More comprehensive information of DEGs from the above three tumor datasets can be found in Supplementary file 1.

**Fig 2.**
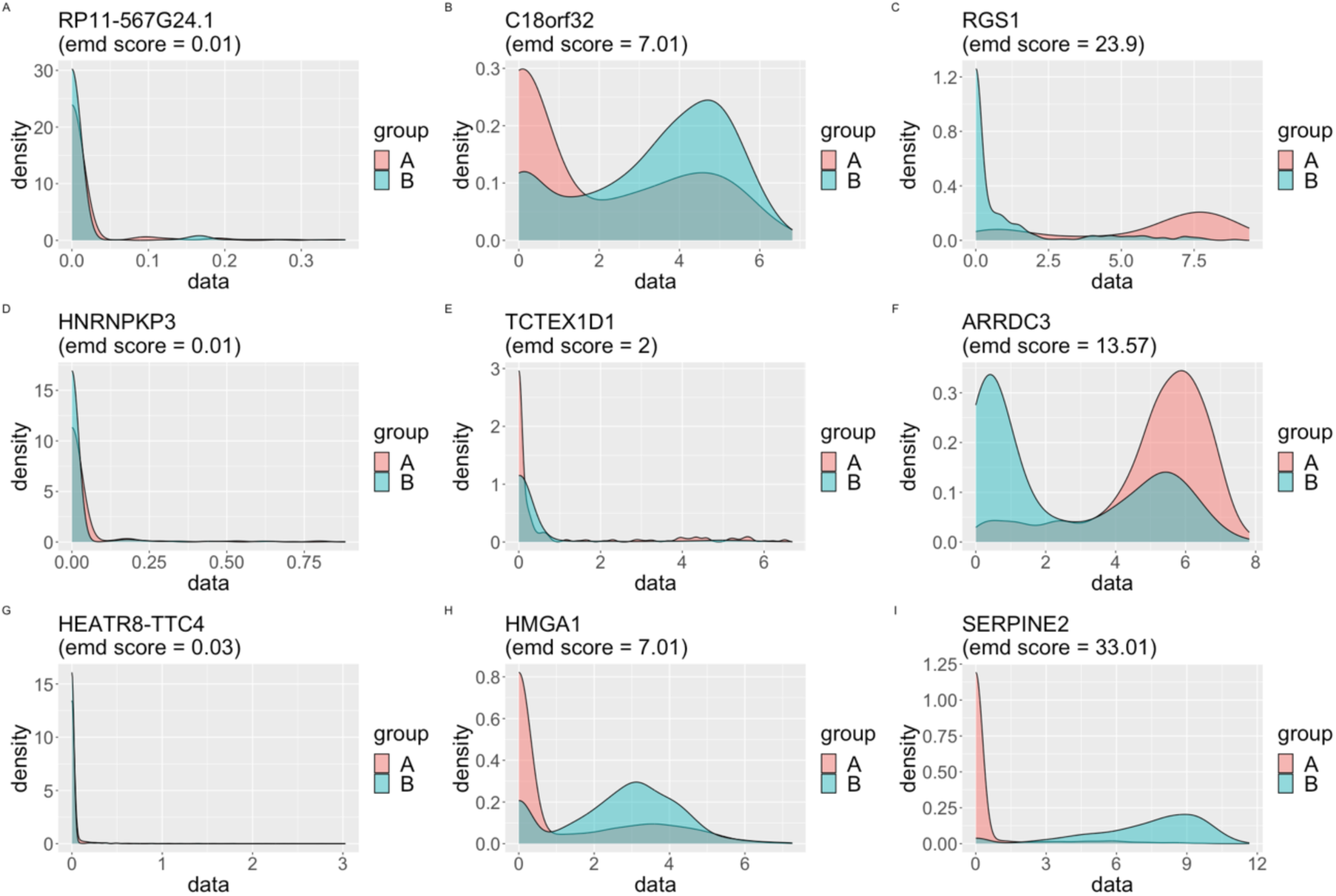
The density plot of all genes using the EMDomics tool between nontumor cells and tumor cells. (A, B, C) The density plots with RP11-567G24.1 (gene), C18orf32 (DEG) and RGS1 (DEG) in the breast cancer dataset. (D, E, F) The density plots with HNRNPKP3 (gene), TCTEX1D1 (DEG) and ARRDC3 (DEG) in the chronic myeloid leukemia dataset. (G, H, I) The density plots with HEATR8-TTC4 (gene), HMGA1 (DEG) and SERPINE2 (DEG) in the melanoma dataset. Group A represents the set of tumor cells. Group B represents the set of nontumor cells.

### Distinguishable performance analysis between tumor and nontumor

To quantitatively evaluate the proposed CSDGI method, we run it between nontumor cells and tumor cells in the breast cancer, chronic myeloid leukemia and melanoma data. For all real tumor scRNA-seq data, we set the parameter about the cluster 𝑘 to 2 in the CSDGI framework to select distinguishable genes between nontumor cells and tumor cells. As shown in Supplementary file 2, we list selected top-20 genes respectively for breast cancer, chronic myeloid leukemia and melanoma data. Here, we perform classification and clustering for selected top-20 genes. To compare different methods, we use Recursive Feature Elimination (RFE) and Chi-square Test (Chi2) as gene identification methods to select genes. For the classification task, we use Support Vector Machine (SVM) and Random Forest (RF) as classification models. For the clustering task, we use Gaussian Mixed Model (GMM) and K-means clustering (K-means) [39] as clustering models. As shown in Fig 3(A, D, G), we find the classification accuracy increases progressively as the number of genes increases for different gene identification methods and classification models. However, the accuracy stabilizes when the number of selected genes fluctuates from 10 and 20. The proposed CSDGI method has outperformed other methods for SVM and RF classification tasks in all cases of selected genes from 2 to 20. Similarly, we also use F1-score to evaluate classification tasks. As shown in Fig 3(B, E, H), our method has still the best classification results than other methods. The reasonable explanation is that the proposed method places its emphasis on identifying and selecting the most relevant genes with specific cell subpopulation. These selected genes may be considered to be the potential driver genes for influencing the characteristics of cancer subtypes. For the clustering tasks, we also compare different methods by using GMM and K-means in the different numbers of selected genes from 2 to 20. As shown in Fig 3(C, F, I), we can find that the clustering ARI of each method has obvious fluctuations with the number of genes from 2 to 8 rather than continuously increasing. Besides, as the increasing number of genes from 8 to 20, the clustering ARI is sometimes reduced instead of rising steadily. However, our method has still outperfprmed other methods in most cases of selected gene numbers from 2 to 20. Therefore, the proposed CSDGI can effectively identify potential driver genes, which helps distinguish different cancer subtypes more correctly. These above classification and clustering results in the breast cancer dataset, chronic myeloid leukemia dataset and melanoma dataset also show that these selected genes have superior distinguishability for characterizing different cell subpopulations.

**Fig 3.**
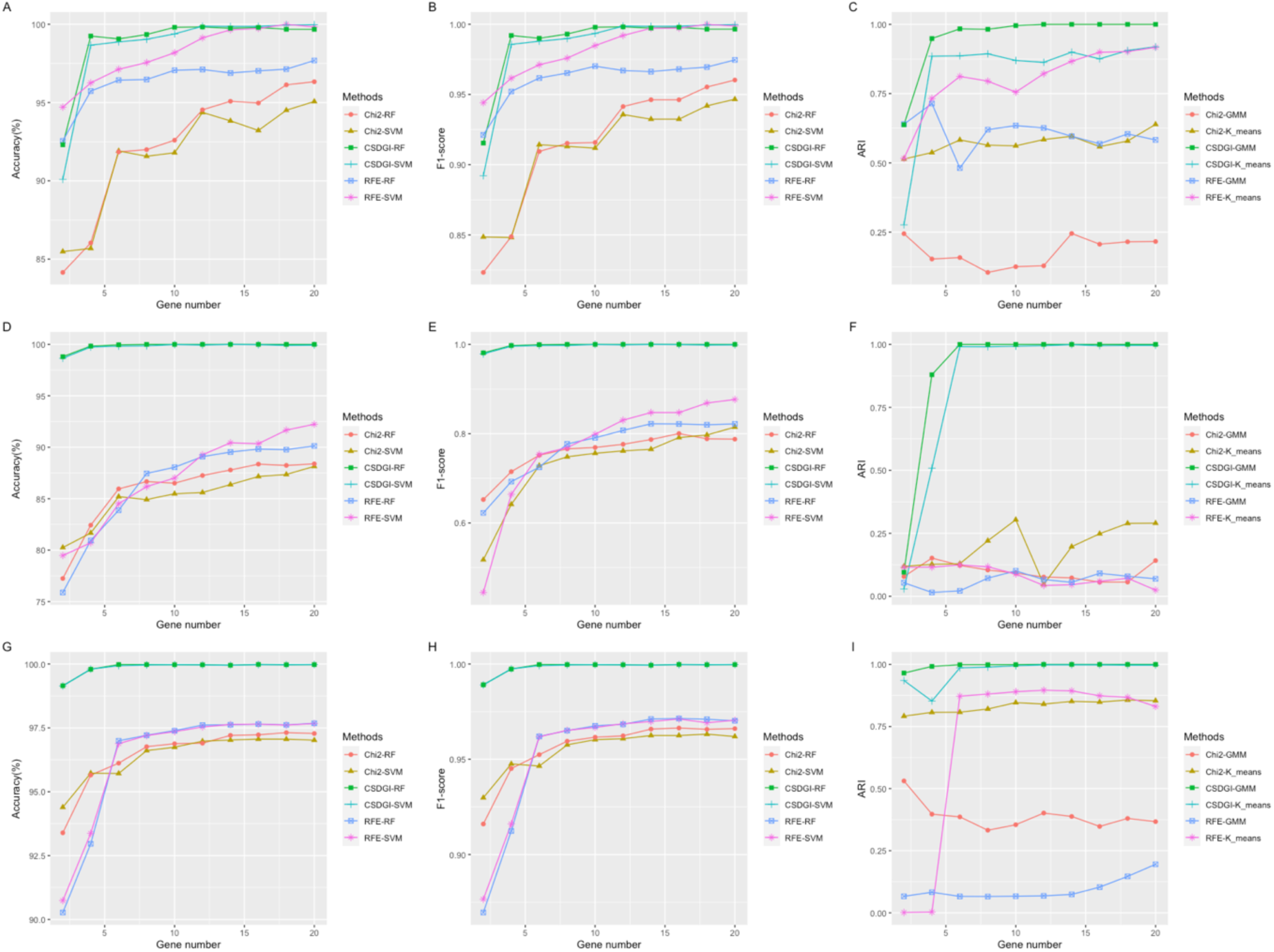
Classification and clustering evaluation results for selected top 20 genes in three real tumor scRNA-seq datasets. (A, B, C) The Accuracy, F1-score and ARI plots in the breast cancer dataset. (D, E, F) The accuracy, F1-score and ARI plots in the chronic myeloid leukemia dataset. (G, H, I) The accuracy, F1-score and ARI plots in the melanoma dataset. For the comparison of different methods, the number of selected genes is 2, 4, 6, 8, 10 12 14, 16, 18 and 20, respectively.

### Estimation of cancer subtypes

To identify cancer subtypes, we apply the proposed CSDGI to the tumor cells in the breast cancer, chronic myeloid leukemia and melanoma data. After excluding nontumor, normal and benign tumor cells, there are 317 breast tumor cells, 902 chronic myeloid leukemia tumor cells and 1257 malignant melanoma cells in these real datasets. Due to the absence of a ground truth of cancer subtypes, we fail to obtain the clustering information of cancer subtypes. Previous studies have acknowledged that it is difficult to determine the number of clusters in a clustering task [39]. To analyze real tumor cells, the sum of squared error (SSE) is employed to identify the number of potential cancer subtypes, where the SSE funnction tool is implemented in the R package (factoextra). As shown in Fig 4(A, C, E), we investigate various cancer subtype number ranging from 1 to 10 by using the Sum of Squared Errors (SSE) method about cluster number identification. Fig 4 (A, C, E) exhibit the elbow point for each tumor cell dataset, which indicates significant changes in the gradient between 1 and 10. Consequently, we select 2, 2, 3 as the optimal value of cancer subtype number, respectively, for the breast cancer tumor cells, chronic myeloid leukemia tumor cells and the malignant melanoma cells. In Fig 4(B, D, F), we also visualize breast tumor cells, chronic myeloid leukemia tumor cells and malignant melanoma cells according to the optimal number. To apply the proposed CSDGI method to three real tumor cell data, we set the parameter 𝑘 to 2, 2, 3, respectively. For these tumor cells from three real cancer datasets, more detailed information about cancer subtypes can be found in Supplementary file 3.

**Fig 4.**
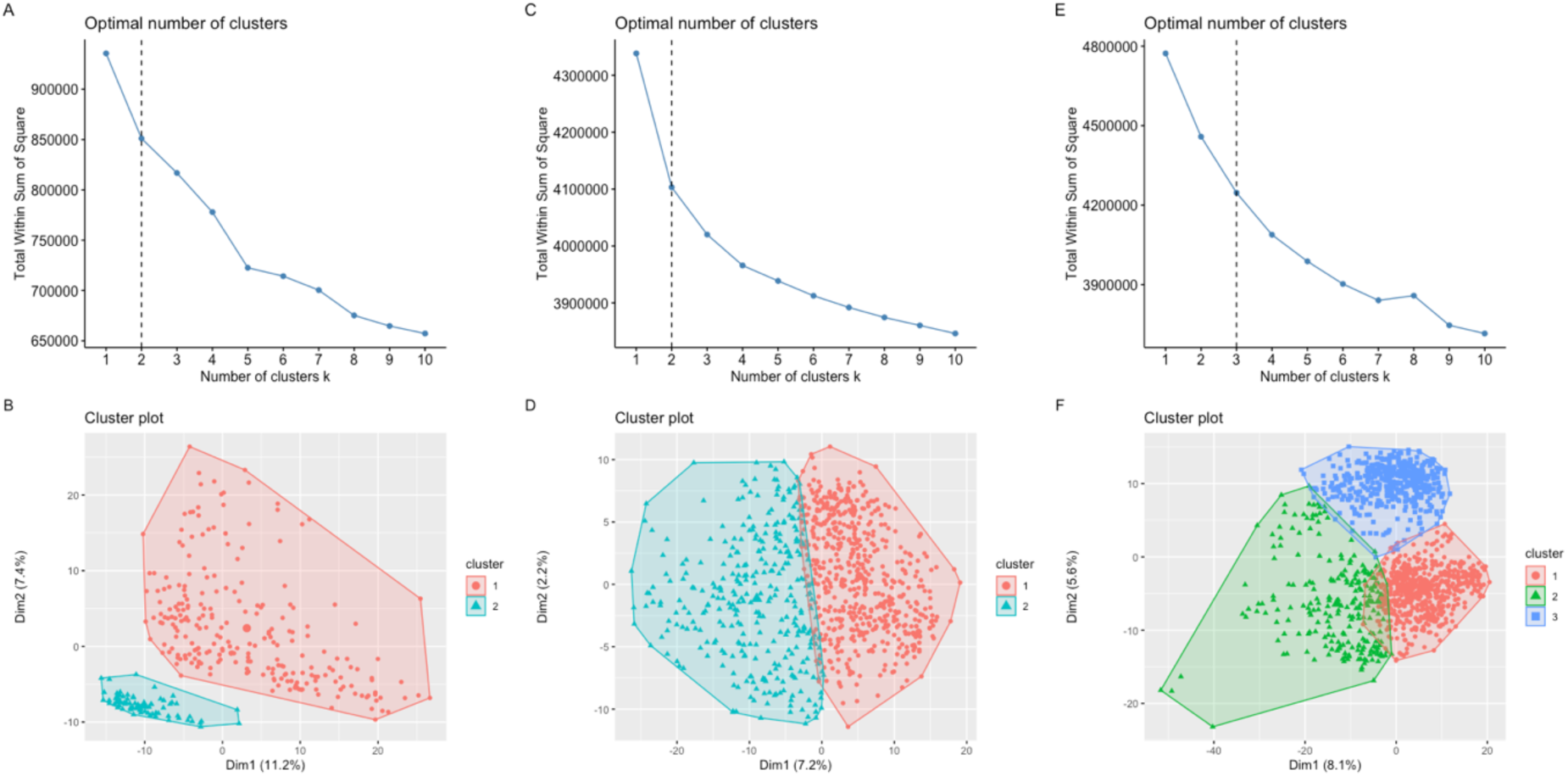
Estimate the number of cancer subtypes for three real tumor cells datasets. (A, B) The optimal value of cancer subtype number and visualization of tumor cells with two possible cancer subtypes in the breast cancer tumor cells. (C, D) The optimal value of cancer subtype number and visualization of tumor cells with two possible cancer subtypes in the chronic myeloid leukemia tumor cells. (E, F) The optimal value of cancer subtype number and visualization of malignant tumor cells with three possible cancer subtypes in the malignant melanoma tumor cells.

### Discovering cancer subtype-specific driver genes

To infer driver genes specific to each cancer subtype, we obtain the learnable low-rank weight in the proposed unsupervised CSDGI framework. Here, we identify top 5% genes as driver genes corresponding to each cancer subtype. As shown in Table 2, we obtain 41 driver genes for each cancer subtype of breast tumor cells, respectively. These underlined genes are genes shared by Subtype 1 and Subtype 2. For the chronic myeloid leukemia tumor cells and the malignant melanoma cells, we infer 52 and 59 driver genes for each cancer subtype, respectively. As shown in Supplementary file 4, we list all detailed information of driver genes. Besides, we also highlight the shared driver genes of each cancer subtype by using the bold in Supplementary file 4.

**Table 2.**
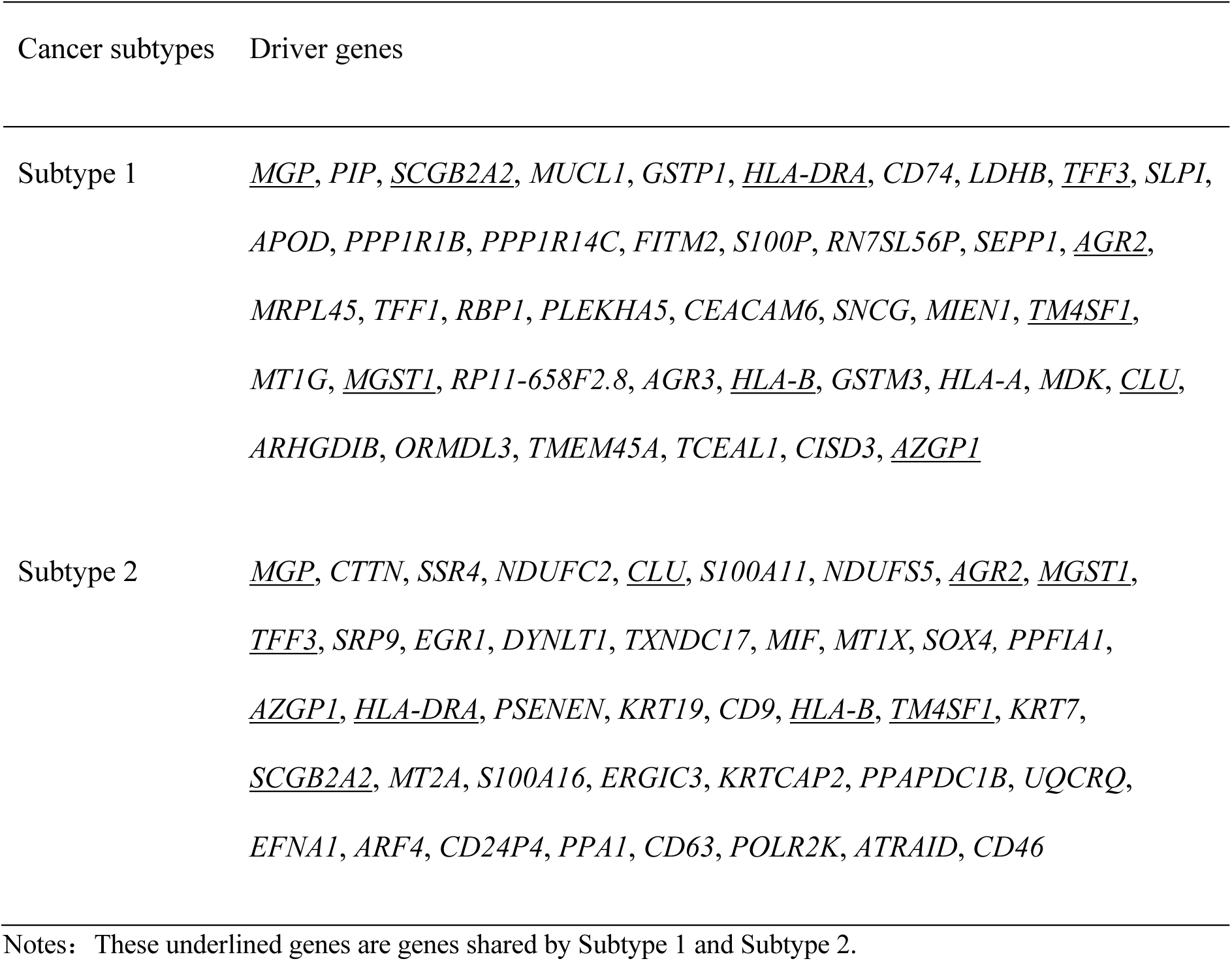
The identified cancer subtype-specific driver genes in the breast tumor cells.

To unveil the biological meaning of driver genes, we perform a more detailed analysis. As shown in Fig 5, we obtain driver gene co-expression network of each cancer subtype by we obtain driver gene co-expression network of each cancer subtype by calculating correlation between driver genes. In Table 2, we can find that *MGP*, *SCGB2A2*, *HLA*, *TFF3*, *AGR2*, *TM4SF1*, *MGST1*, *HLA-B*, *CLU* and *AZGP1* are shared in two subtypes for the breast tumor cells. Previous studies show that *MGP* promotes the breast cancer proliferation [40], *SCGB2A2* and *TFF3* can be viewed a valuable predictive biomarker in breast cancer [41, 42]. *MUCL1* are identified as driver genes in subtype 1, which have been found to be critical markers of breast cancer [43]. In Fig 5(A, B), *MGP*, *SCGB2A2* and *TFF3* have been marked with red box. For the chronic myeloid leukemia tumor cells, the shared driver genes between subtype 1 and 2 include *RPL32*, *SAT1*, *SNORD102*, *CD52*, *TSTD1*, *SELL*, *ATRAID*, *PSMB6*, *NFKBIA*, *NDUFA12* and *DBI* in Supplementary file 4. Previous evidences illustrate that *RPL32* plays an important role in the SF3B1 mutation for chronic myeloid leukemia (CML) disease progression [44], *CD52* helps identify molecular signature of CML [45] and *NFKBIA* plays a strategic role in CML molecular response [46]. In addition, *LGALS1* is identified as driver genes in the CML cancer subtype 1 and previous works also find that *LGALS1* can be selected as a critical biomarker of CML [47]. In Fig 5(C, D), *RPL32*, *CD52*, *NFKBIA* and *LGALS1* have been marked with red box. For the malignant melanoma cells, *PMEL*, *IFITM1*, *HSPA1A*, *DUSP1*, *FOS* and *TMSB4X* are the shared driver genes across three subtypes in Supplementary file 4. Previous studies have indicated that *PMEL* is a important link between Parkinson’s disease and melanoma [48]. Other researchers have also regarded it as a prognostic predictor in skin cutaneous melanoma (SKCM) [49]. Furthermore, the shared *RGS* between Subtype 1 and Subtype 3, and the shared *MIA* between Subtype 1 and Subtype 2 have been viewed as important biomarkers in malignant melanoma [50, 51]. In Fig 5(E, F, G), *PMEL*, *RGS1*and *MIA* have been marked with red box. Overall, these cancer subtype-specific driver genes indicates more significant biological meaning to study the breast tumor cells, chronic myeloid leukemia tumor cells and malignant melanoma cells.

**Fig 5.**
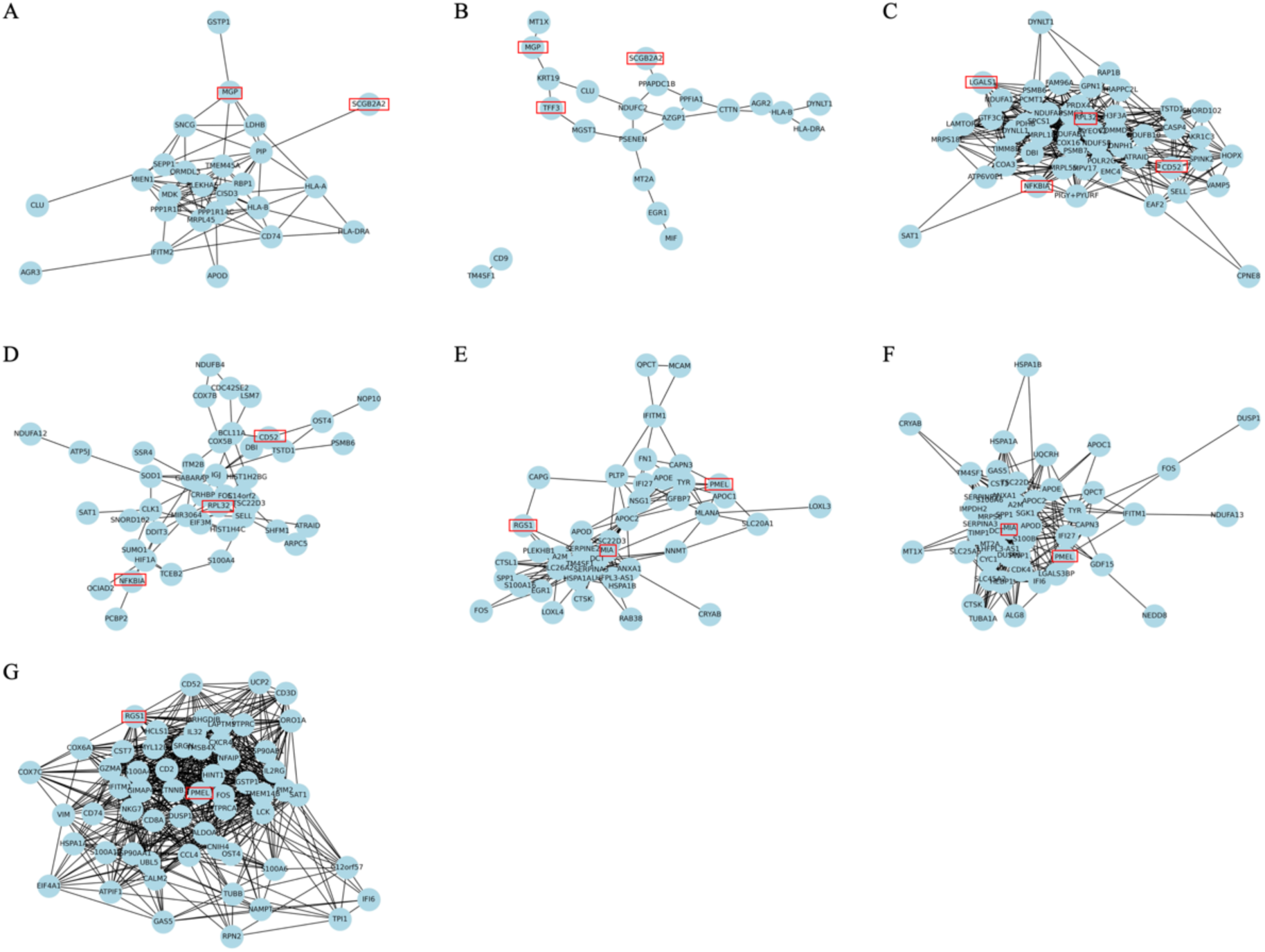
Driver gene co-expression network of each cancer subtype for three real tumor cells datasets. (A, B) Cancer subtype 1 and 2 in the breast cancer tumor cells. (C, D) Cancer subtype 1 and 2 in the chronic myeloid leukemia tumor cells. (E, F, G) Cancer subtype 1, 2 and 3 in the malignant melanoma tumor cells.

### Functional and disease enrichment analysis for CSDGs

To investigate the potential biological significance of CSDGs inferred by our CSDGI method, we perform the functional and disease enrichment analysis. We use Gene Ontology (GO, http://www.geneontology.org/) to analyze the biological function: biological process (BP), cellular component (CC), and molecular function (MF). Besides, the Kyoto Encyclopedia of Genes and Genomes Pathway (KEGG, http://www.genome.jp/kegg/) and Reactome Pathway (Reactome, http://reactome.org/) are also included in functioanal enrichment analysis. For the disease enrichment analysis, we use Disease Ontology (DO, http://disease-ontology.org/), DisGeNET (http://www.disgenet.org/) and Network of Cancer Genes database (NCG, http://ncg.kcl.ac.uk/) to analyze the inferred driver genes of each cancer subtype. As a result, we set the threshold of significance enrichment *p-value* to 0.01. As shown in Supplementary file 5, we list comprehensive information on functional and disease enrichment analysis about CSDGs for the breast cancer tumor cells, the chronic myeloid leukemia tumor cells and the malignant melanoma cells. Previous studies show that cell aggregation and mineral absorption have a significant influence on the progression of breast cancer [52, 53]. Polyamine biosynthetic process promotes CML tumor growth [54] and ferroptosis may be a novel strategy for chronic myeloid leukemia anti-tumor therapy [55]. Cell aggregation, mineral absorption, polyamine biosynthetic process, and ferroptosis can be viewed as important enrichment analysis results in in Supplementary file 5. For the malignant melanoma cells, the enrichment analysis also indicates more important melanoma-related biological processes including melanin biosynthetic process, melanin metabolic process and melanosome organization in Supplementary file 5. Furthermore, we find that genes enriched on the same term type are different driver genes of different cancer subtypes for a group of tumor cells. This may be because the informative CSDGs exhibit differences between cancer subtypes. All in all, these inferred CSDGs can serve as valuable references to drive cancer subtype growth and finding new tumor driver markers, which illustrates the biological meaning of tumor development and helps understand the mechanisms of cell transformation driving tumours.

## Discussion and conclusions

Cancer is a heterogeneous disease, where cancer driver genes can drive tumorigenesis and the unstable cellular growth. CDGs show the characteristics of tissue-specific or condition-specific. Thus, to deeply understand the cancer progression mechanisam at the cancer subtype level, we need to uncover CSDGs. However, most of the existing computational methods have mainly used the cohort information rather than the cancer subtype-specific information to infer CDGs. Tumors of different subtype are highly heterogeneous. Therefore, we need to identify CDGs from tumor cells of each cancer subtype. In this work, we use the unsupervised learning mechanism to propose a novel CSDGI computational method. At single-cell level, we use CSDGI to infer CSDGs by only considering single-cell transcriptomics data. In the proposed CSDGI, the Encoder-Decoder-Framework helps identify potential cancer subtypes. Furthermore, the application to the breast cancer tumor cells, the chronic myeloid leukemia tumor cells and the malignant melanoma tumor cells indicates that CSDGI may be a new solution to explain cancer subtype molecular mechnism. One possible limitation of the current CSDGI framework is that the inferred CSDGs relys on the result of cancer subtype identification. The different tumor cells in the same cancer subtype may result in different CSDGs. Another limitation is that the current CSDGI method may be better suited to identify CDGs in the near 1000-dimension gene space. For the high-dimension gene expression data, such as near 20,000-dimension gene space without gene filtering, the proposed CSDGI could not infer CSDGs more efficiently. In summary, we believe that CSDGI is fundamental to explore specific biological function involved in tumorigenesis across cancer subtypes, and improve existing driver gene identification methods. The cancer subtype-specific framework may provide a new solution to deeply understand the mechanisms of cell transformation and cancer progression, and give a shortlist of biologically meaningful genes that have potential to promote precision cancer medicine.

## Supporting information

Supplementary file 1: Differential expression genes in three scRNA-seq datasets.

Supplementary file 2: Selected top 20 genes in three scRNA-seq datasets.

Supplementary file 3: Identified cancer subtypes in three scRNA-seq datasets.

Supplementary file 4: The identified cancer subtype-specific driver genes in the chronic myeloid leukemia tumor cells and the malignant melanoma cells.

Supplementary file 5: Functional enrichment analysis and disease enrichment analysis in three scRNA-seq datasets.

## Supplementary Data

Supplementary data are available online.

## Data availability statement

The source code used to replicate all our analyses, including all real data, is available at the following link: https://github.com/linxi159/CSDGI.

## Acknowledgments

We thank the editor and reviewers for their help and comments during the preparation of the manuscript.

## Author contributions

Conceptualization: Xiucai Ye.

Data curation: Meng Huang.

Formal analysis: Meng Huang, Jiangtao Ma, Guangqi An, Xiucai Ye.

Investigation: Meng Huang, Jiangtao Ma.

Methodology: Meng Huang, Jiangtao Ma, Xiucai Ye.

Project administration: Xiucai Ye.

Resources: Guangqi An, Xiucai Ye.

Software: Meng Huang, Jiangtao Ma.

Supervision: Xiucai Ye.

Validation: Meng Huang.

Visualization: Meng Huang, Guangqi An.

Writing – original draft: Meng Huang, Jiangtao Ma.

Writing – review & editing: Meng Huang, Jiangtao Ma, Xiucai Ye.

